# Cortical organoids from congenital DM1 PSCs reveal MBNL-dependent corticogenesis defects and enable preclinical testing of therapeutic compounds

**DOI:** 10.64898/2026.07.23.740263

**Authors:** Azania Abatan, Jérôme Polentes, Margot Bouquier, Adeline Beuriot, Olivier Chose, Hamel Mahiou, Lina El Kassar, Karine Giraud-Triboult, Laure Chatrousse, Alexandra Benchoua, Stéphanie Tomé, Geneviève Gourdon, Mario Gomes-Pereira, Sandrine Baghdoyan, Cécile Martinat

## Abstract

Myotonic dystrophy type 1 (DM1) is caused by an expansion of a CTG repeat in the 3′ untranslated region of the *DMPK* gene, leading to accumulation of toxic CUG-repeat RNAs, sequestration of MBNL proteins and widespread splicing dysregulation. Congenital DM1 (CDM), the most severe form of the disease, is associated with profound muscular and neurodevelopmental defects, yet the mechanisms underlying early human brain involvement remain poorly understood. Here, we generated cortical organoids from patient-derived pluripotent stem cells carrying >1000 CTG repeats, an expansion typically associated with CDM, to model early human neurodevelopment. DM1 molecular and cellular hallmarks were detected at early developmental stages, including nuclear *DMPK* RNA foci in neural progenitor cells and reduced proliferative capacity. As organoids matured, CDM cultures displayed altered cortical composition, with reduced CTIP2⁺ and SATB2⁺ neuronal populations and increased NFIA⁺/GFAP⁺ glial cells. In parallel, 120-day-old organoids recapitulated splicing abnormalities previously identified in DM1 patient brain tissue. To assess the contribution of MBNL dysfunction, we analyzed cortical organoids derived from *MBNL2* and *MBNL1/2/3* knockout induced pluripotent stem cells, which reproduced key neurodevelopmental phenotypes observed in CDM organoids, supporting a central role for MBNL loss of function in impaired corticogenesis. Finally, we evaluated the translational relevance of this model using tideglusib and erythromycin, two compounds currently under clinical evaluation in DM1 patients. Both treatments reduced *DMPK* RNA foci and restored proliferation defects in SOX2⁺ neural progenitors. Together, these findings establish cortical organoids as a robust human model of CDM-associated neurodevelopmental defects, uncover MBNL-dependent mechanisms underlying early corticogenesis impairment and demonstrate the utility of this platform for translational therapeutic discovery in DM1.

## INTRODUCTION

Myotonic dystrophy type 1 (DM1) is a dominantly inherited multisystemic disorder caused by CTG repeat expansion in the 3′ untranslated region of the *DMPK* gene. The resulting toxic CUG RNA sequesters MBNL proteins and disrupts alternative splicing, establishing DM1 as a prototypical spliceopathy (**Mankodi et al., 2001; Miller et al., 2000; Kuyumcu-Martinez et al., 2007; Hicks et al., 2024)**. Disease severity increases with repeat length, culminating in congenital DM1 (CDM), the most severe form of the disease (**Harley et al., 1993**). CDM patients present at birth with severe muscular hypotonia together with cognitive, behavioral and adaptive impairments, pointing to early neurodevelopmental dysfunction (**Ho et al., 2015; De Antonio et al., 2016**). However, despite growing evidence for central nervous system involvement, therapeutic efforts have predominantly targeted skeletal muscle pathology. Neuroimaging and neuropathology studies further reveal cortical abnormalities, including white matter defects, ventricular enlargement, and cortical atrophy, suggesting impaired corticogenesis as a primary pathogenic mechanism (**Krieger et al., 2024; Wozniak et al., 2011; Shear et al., 2024**).

Progress in understanding CNS pathology and evaluating potential therapies has been hindered by the absence of human models that faithfully recapitulate early cortical development. Animal models and 2D cellular systems capture RNA gain-of-function mechanisms, but they fail to recapitulate the spatiotemporal complexity underlying the emergence and organization of neural and glial progenitor populations during human corticogenesis (**Lee et al., 2019; Xia, et al., 2013; Denis et al., 2013; Eltahir et al., 2022**). Human iPSC-derived cortical organoids provide a transformative platform, recapitulating neural progenitor expansion, neuronal specification, and glial differentiation while reproducing DM1 molecular hallmarks such as RNA foci, MBNL sequestration, and splicing defects (**Morelli et al., 2022; De Serres-Berard, 2025**).

In this study, we generated cortical organoids from CDM patient-derived iPSCs and MBNL- depleted lines to investigate how toxic CUG RNA impairs early corticogenesis and neuroglial development. Importantly, we explored the potential of these organoids as a preclinical platform to evaluate clinically investigated compounds targeting CNS manifestations often overlooked in DM1 interventions. This approach establishes a human-relevant system that bridges mechanistic insight and translational application, providing a framework to extend therapeutic evaluation beyond muscle and toward the neurodevelopmental component of DM1.

## RESULTS

### Generation of CDM cortical organoids

To investigate early corticogenesis in congenital DM1, three DM1 hPSC lines harboring more than 1300 CTG repeats (CDM) and three human control lines (CTL) were differentiated into cortical organoids using a stepwise protocol (**Paşca et al., 2015**) (**Figure 1A**). At days 30 and 60 of differentiation, immunostaining revealed abundant SOX2⁺/PAX6⁺ neural progenitor cells and HuC/D⁺/TUJ1⁺ postmitotic neurons in organoids derived from both control and CDM lines, indicating preserved early cortical patterning and lineage specification (**Figure 1B**). Extension of differentiation to 120 days enabled the detection of GFAP⁺ and S100β⁺ astrocytic cells in both groups, demonstrating that the neurogenic-to-gliogenic transition occurred in CDM as well as control organoids (**Figure 1B**). To confirm the persistence of the pathogenic RNA hallmark, RNA fluorescence in situ hybridization was performed throughout differentiation. Prominent intranuclear CUG RNA foci were detected in approximately 70% of nuclei in CDM organoids at both days 30 and 120 (**Figure 1C**), indicating sustained expression of the mutant *DMPK* transcript. Together, these findings establish CDM cortical organoids as a robust human model to investigate the cellular and molecular consequences of toxic RNA expression during early corticogenesis.

**Figure 1.**
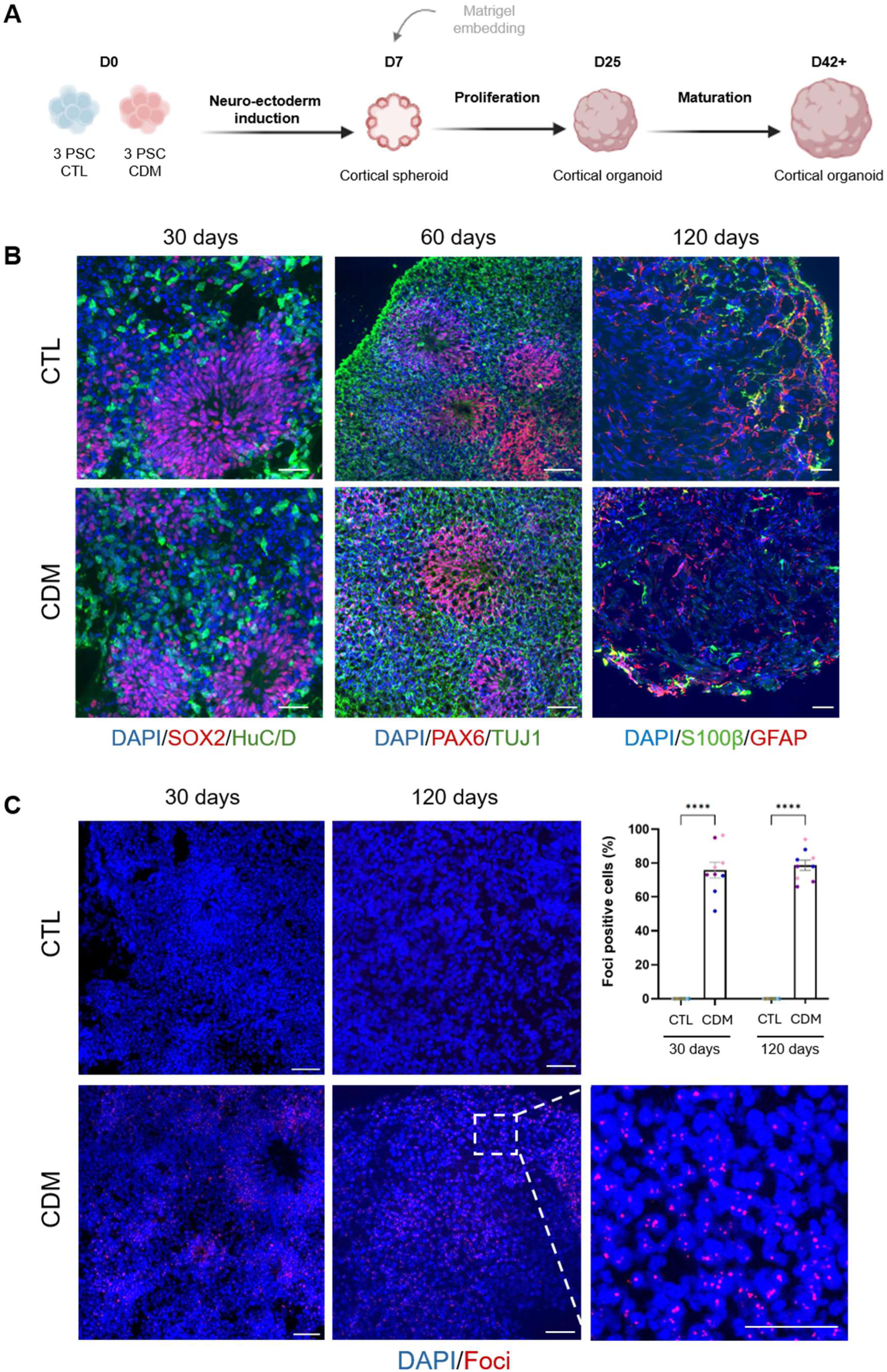
Generation of cortical organoids derived from CDM and control human pluripotent stem cells. **(A)** Schematic representation of the protocol used to generate control (CTL) and CDM cortical organoids. **(B)** Immunostaining of SOX2, PAX6, HuC/D, TUJ1, S100 β and GFAP markers in control and CDM cortical organoids at days 30, 60 and 120 of differentiation **(C)** Representative images and quantification of RNA-FISH showing foci in CTL and CDM cortical organoids at days 30 and 120 of differentiation. Colors indicate distinct cell lines within each genotype, and individual dots represent the mean value of three organoids from one independent differentiation experiment. Data are presented as mean ± SD. Statistical analysis was performed using two-way ANOVA followed by Dunnett’s multiple comparisons test. Statistical significance is indicated as follows: ****p < 0.0001. Scale bars, 50 µm.

### RNA foci accumulation and MBNL-dependent splicing defects during early cortical differentiation

The early detection of *DMPK* RNA foci in CDM organoids prompted us to examine their distribution across neural cell populations. RNA-FISH combined with immunostaining showed no significant difference in the proportion of foci-positive cells between SOX2^+^ progenitors and HuC/D^+^ neurons. However, the number of foci per cell was significantly higher in progenitor cells, indicating a greater RNA burden at early developmental stages (**Figure 2A- C**). Both MBNL1 and MBNL2 are expressed during fetal development, with MBNL2 gradually becoming predominant in the adult brain (**Taylor et al., 2023**). The analysis by RT-qPCR of *DMPK*, *MBNL1* and *MBNL2* transcripts during cortical organoids differentiation confirmed an increase of expression of *DMPK* and *MBNL2* genes that becomes statistically significant at day 60 while *MBNL1* transcript expression remained stable over time (**Figure 2D**). Sequestration of MBNL2 within nuclear *DMPK* RNA foci was observed by combined RNA-FISH and immunofluorescence in 30-day-old CDM organoids, suggesting that functional depletion occurred at early stages of differentiation (**Figure 2E**). The presence of the mutated *DMPK* transcript in 120-day-old CDM organoid was associated with reduced levels of MBNL2 protein expression detected by western blot analysis compared to CTLs (**Figure 2F**). To assess the functional impact of MBNL1/2 depletion, alternative splicing events known to be dysregulated in DM1 brain tissue, including *MBNL2* exon 5, *ATP2A1* exon 22, and *ITGA6* exon 27 (**Otero et al., 2021**), were analyzed throughout differentiation. Although modest changes in splicing were observed at days 30 and 60, none reached statistical significance. By day 120, control organoids exhibited a transition in splicing profiles that was not detected in CDM or *MBNL*- knockout cortical organoids, indicating that toxic RNA and/or MBNL loss may interfere with the molecular processes underlying differentiation. (**Figure 2 G-I, Figure S1**). These defects were comparable in timing and magnitude to those detected in cortical organoids derived from *MBNL2* knockout and *MBNL1*/2/3 triple knockout hiPSCs supporting that MBNL2 sequestration in *DMPK* foci contributes to splicing defects of 120-day-old CDM cortical organoids (**Figure 2, Figure S1**). Collectively, these results demonstrate that *DMPK* RNA foci affect both neural progenitors and neurons in CDM organoids. Their presence is associated with MBNL mislocalization and MBNL-dependent splicing differences that emerge over time and become detectable by day 120.

**Figure 2.**
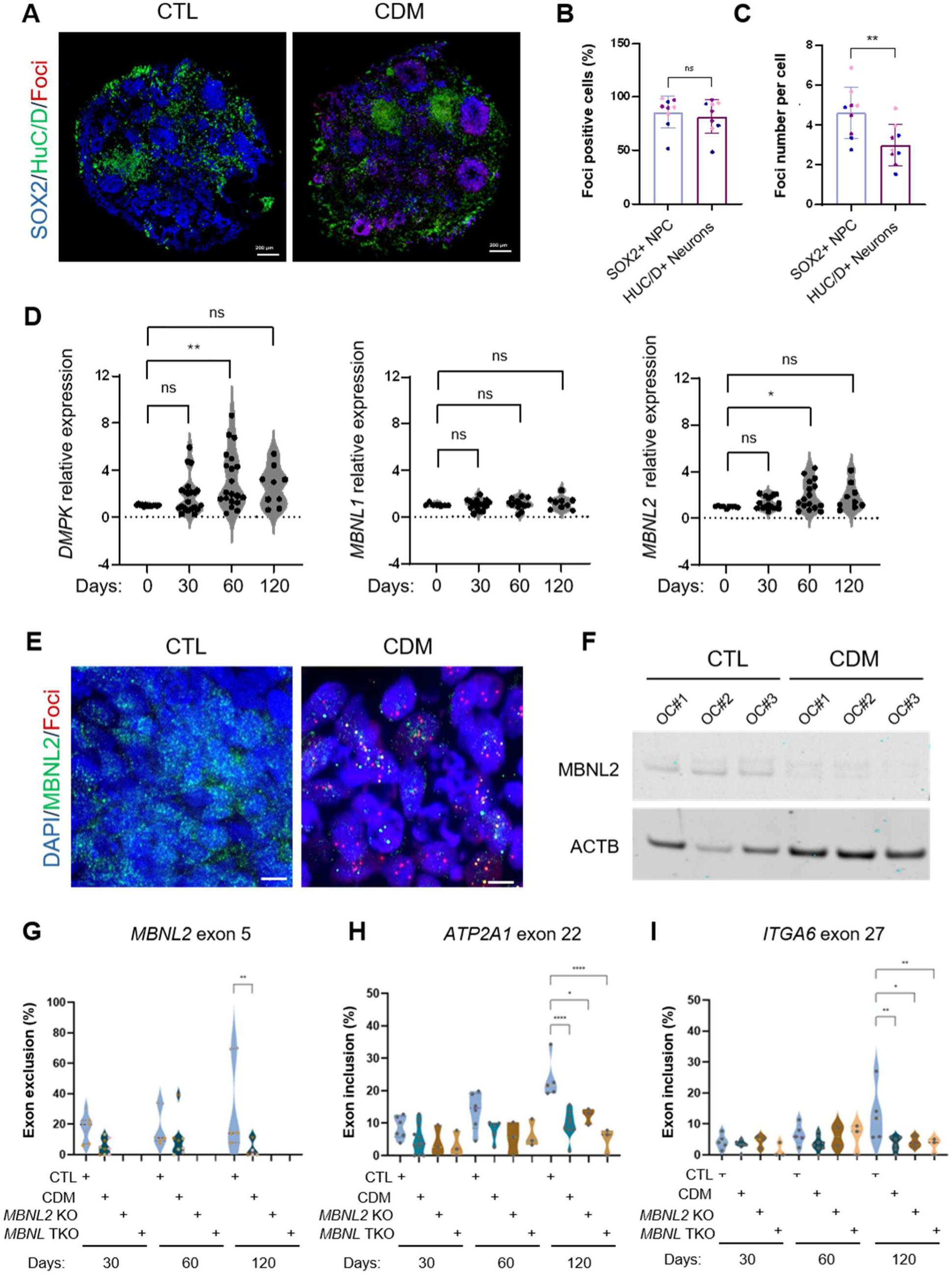
Detection of DM1 molecular hallmarks in CDM and *MBNLs* KO cortical organoids. (**A-C**) Representative of RNA-FISH images and quantifications showing foci in SOX2^+^ neural progenitors (NPC) and HuC/D^+^ neurons in 30-day-old CTL and CDM cortical organoids. Scale bars 200 µM. Colors indicate distinct cell lines within each genotype, and individual dots represent the mean value of three organoids from one independent differentiation experiment. (**D**) RT-qPCR analysis of *DMPK*, *MBNL1* and *MBNL2* gene expressions in CTL and CDM cells at day 0 and in cortical organoids at days 30, 60, and 120 of differentiation. (**E**) Immunodetection of MBNL2 sequestration in RNA foci by RNA-FISH in 30-day-old CDM cortical organoids. Scale bars, 10 µm. (**F**) Representative western blot analysis of MBNL2 and ACTB expressions in 120-day-old in three batches of CTL and CDM cortical organoids (OC#1 to #3). (**G-I**) Quantitative analysis of alternative splicing for *MBNL2*, *ATP2A1* and *ITGA6* transcripts by RT-PCR in 30-, 60- and 120-day-old cortical organoids. Colors indicate distinct cell lines within each genotype, and individual dots represent the mean value of three organoids from one independent differentiation experiment. Data are presented as mean ± SD. Statistical analyses were performed using unpaired two-tailed Student’s t-test and one-way ANOVA followed by Dunnett’s multiple comparisons test. Statistical significance is indicated as follows: *p < 0.05; **p < 0.01; ****p < 0.0001.

### Reduced proliferation of cortical neural progenitors in CDM and MBNL-depleted organoids at early differentiation stages

Given the higher number of *DMPK* RNA foci observed in progenitors, their effect on the neural progenitor cell (NPC) pool was examined. Immunostaining of SOX2^+^ NPCs in 30-days-old organoids revealed a significant reduction in progenitor numbers in CDM organoids relative to controls. A similar decrease was also observed in *MBNL2* KO and *MBNL* TKO organoids suggesting a contribution of MBNL depletion to this phenotype (**Figure 3A-B**). Apoptotic activity, assessed by cleaved Caspase 3 staining in SOX2⁺ progenitors, was similar across CDM, *MBNL2* KO, *MBNL* TKO, and control organoids at day 30, suggesting that the reduction in progenitor numbers likely reflects impaired proliferation rather than increased cell death (**Figure 3C-D**). The proliferative capacity of NPCs was then evaluated using EdU incorporation. A significant decrease in EdU⁺ progenitors was detected in CDM, *MBNL2* KO, and *MBNL* TKO organoids compared to controls (**Figure 3E-F**). Consistently, the proportion of PHH3⁺ mitotic cells was also reduced in CDM and *MBNL*-depleted organoids (**Figure S2**). These results indicate that impaired proliferation is a predominant contributor to reduced NPC number at days 30 in CDM organoids. Moreover, the similarity of the phenotype in MBNL-depleted organoids underscores the critical role of MBNL proteins in maintaining progenitor expansion during early cortical differentiation.

**Figure 3.**
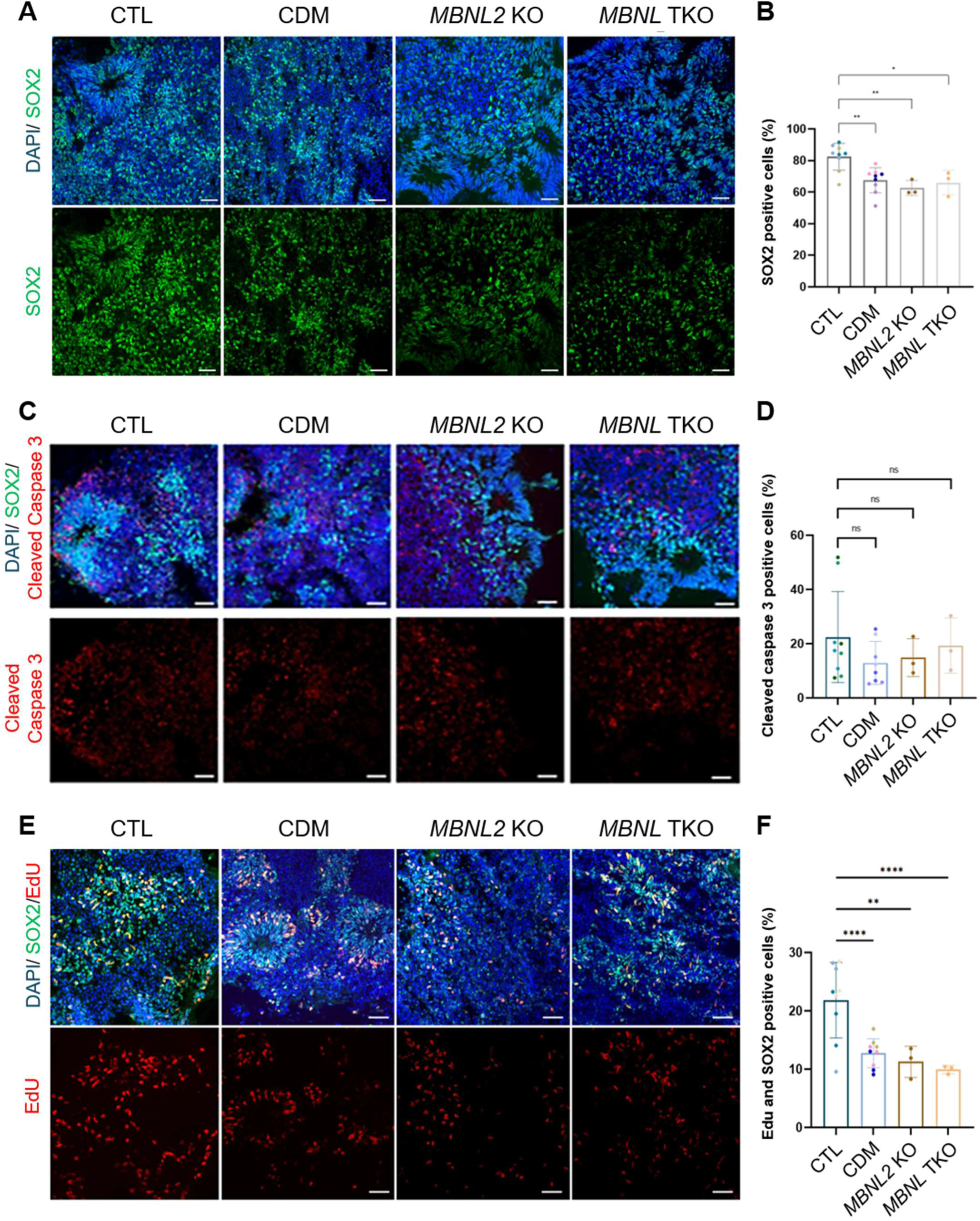
Proliferation defects of SOX2^+^ neural progenitor cells from 30-day-old CDM and *MBNLs* KO cortical organoids. (**A-B**) Representative images and quantification of neural progenitors expressing the SOX2 marker labelled by immunostaining in 30 days-old cortical organoids. (**C-D**) Representative images and quantification of cleaved-caspase 3^+^ SOX2^+^ neural progenitors detected by immunostaining in 30 days-old cortical organoids. Scale bar 50µm. (**E- F**) Representative images and quantification of SOX2^+^ neural progenitors immunostained with EdU. Colors indicate distinct cell lines within each genotype, and individual dots represent the mean value of three organoids from one independent differentiation experiment. Data are presented as mean ± SD. Statistical analysis was performed using two-way ANOVA followed by Dunnett’s multiple comparisons test. Statistical significance is indicated as follows: *p < 0.05; **p < 0.01; ****p < 0.0001. Scale bars, 50 µm.

### Altered neuronal and glial fate specification in 120 days-old CDM and MBNL-depleted organoids

The presence of *DMPK* foci and their effect on corticoid differentiation were next evaluated at later stages. Previous studies have reported an imbalance between neural and glial populations in DM1-derived cortical organoids (**Morelli et al., 2022**). Immunostaining revealed reductions in CTIP2⁺ deep-layer and SATB2⁺ superficial-layer neurons in CDM, *MBNL2* KO, and *MBNL1/2/3* TKO organoids compared to controls (**Figure 4A-D**). Concurrently, GFAP⁺ glial populations and NFI-A⁺ progenitors were increased, indicating enhanced glial specification (**Figure 4E-H**). Analysis of RNA foci in 120-day-old organoids revealed a higher proportion of foci-positive cells among NFIA⁺ cells compared with HuC/D⁺ cells, which also contained fewer foci (**Figure 4I–K**).

**Figure 4.**
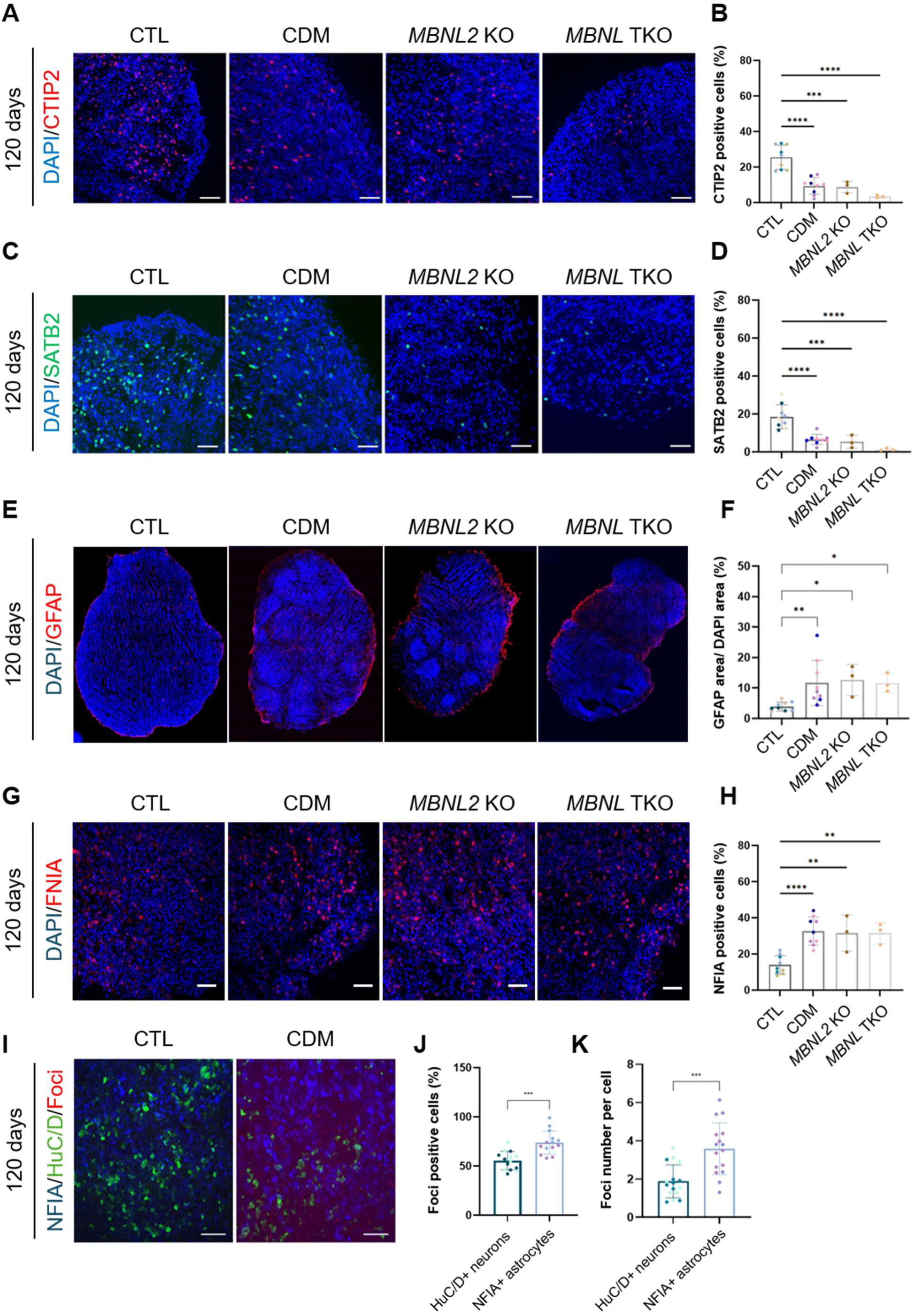
Neuronal and glial defects in CDM and *MBNL* KO 120 days-old cortical organoids. (A-B) Representative images and quantification of CTIP2^+^ neurons in 120-day-old cortical organoids. Scale bars, 50 µm. **(C-D)** Representative images and quantification of SATB2^+^ neurons in 120-day-old cortical organoids. Scale bar, 50 µm. **(E-F)** Representative images and quantification of GFAP^+^ astrocytes in 120-day-old organoids. Scale bars, 200 µm. **(G-H)** Representative images and quantification of NFIA1^+^ astrocytes in 120-day-old organoids. Scale bars, 50 µm. **(I)** Representative images of RNA-FISH showing foci in NFIA1^+^ astrocytes and HUC/D^+^ neurons in CDM 120-day-old organoids. Scale bars, 50 µm. **(J-K)** Quantification of Foci-positive cells and number of foci in CDM astrocytes and neurons. Colors indicate distinct cell lines within each genotype, and individual dots represent the mean value of three organoids from one independent differentiation experiment. Data are presented as mean ± SD. Statistical analysis was performed using one-way ANOVA followed by Dunnett’s multiple comparisons test, a Kruskal–Wallis test followed by Dunn’s multiple comparisons test and an unpaired two-tailed Student’s t-test. Statistical significance is indicated as follows: *p < 0.05; **p < 0.01; ***p < 0.001; ****p < 0.0001.

To further characterize astrocyte-lineage cells and assess the distribution of *DMPK* RNA foci within this population, 120-day-old cortical organoids were dissociated and cells expressing the glutamate aspartate transporter GLAST were purified using magnetic-activated cell sorting. Immunostaining confirmed astrocytic identity, with ∼60% of GLAST⁺ cells expressing GFAP and 80% S100β (**Figure S3A**). RNA-FISH revealed that, while the proportion of foci-positive cells was similar in GLAST⁺ and GLAST⁻ fractions, the number of foci per nucleus was significantly higher in GLAST⁺ cells from CDM organoids (**FigureS3B-E**), consistent with the previous report of higher RNA foci burden in astrocytes in DM1 mouse models (**Dinca et al., 2022**). Together, these results demonstrate a shift from neuronal to glial fate in CDM and MBNL-depleted organoids, suggesting that MBNL proteins contribute to proper glial lineage induction and expansion.

### Drug treatment reveals rescue of DM1 molecular and cellular defects in cortical organoids

Several clinical trials are currently underway for DM1, primarily targeting muscle pathology, as most therapeutic strategies to date have been developed based on muscle phenotypes. The impact of these therapies on CNS defects remains largely unexplored. To evaluate whether CDM cortical organoids could serve as a platform to test CNS-targeted interventions, two compounds with completed phase II trials, Tideglusib (AMO-02) and erythromycin, were assessed (**Horrigan et al., 2020; Nakamori et al., 2024**). Tideglusib, a GSK3β inhibitor, was selected for its reported cognitive benefits in congenital-onset patients (REACH-CDM trial; NCT05004129), whereas erythromycin was chosen for its ability to displace MBNL proteins from *DMPK* RNA foci (**Nakamori et al. 2015**) (jRCT2051190069).

Cortical organoids derived from two CDM iPSC lines were treated from day 10 to day 25 and phenotypes were analyzed at day 30 (**Figure 5A**). Tideglusib significantly reduced *DMPK* transcript levels compared to untreated organoids, whereas erythromycin had no effect on transcript abundance as previously observed in murine C2C12 myoblasts (**Nakamori et al. 2015**) (**Figure 5B**). Both compounds, however, markedly decreased the number of nuclear *DMPK* RNA foci (**Figure 5C-D**). Reduction of RNA foci was accompanied by a rescue of the proliferation deficit. The proportion of EdU⁺ SOX2⁺ neural progenitors increased significantly following treatment with either compound (**Figure 5E-F**). No increase in cleaved caspase-3- positive cells was observed, indicating absence of overt toxicity (**Figure 5G-H**).

**Figure 5.**
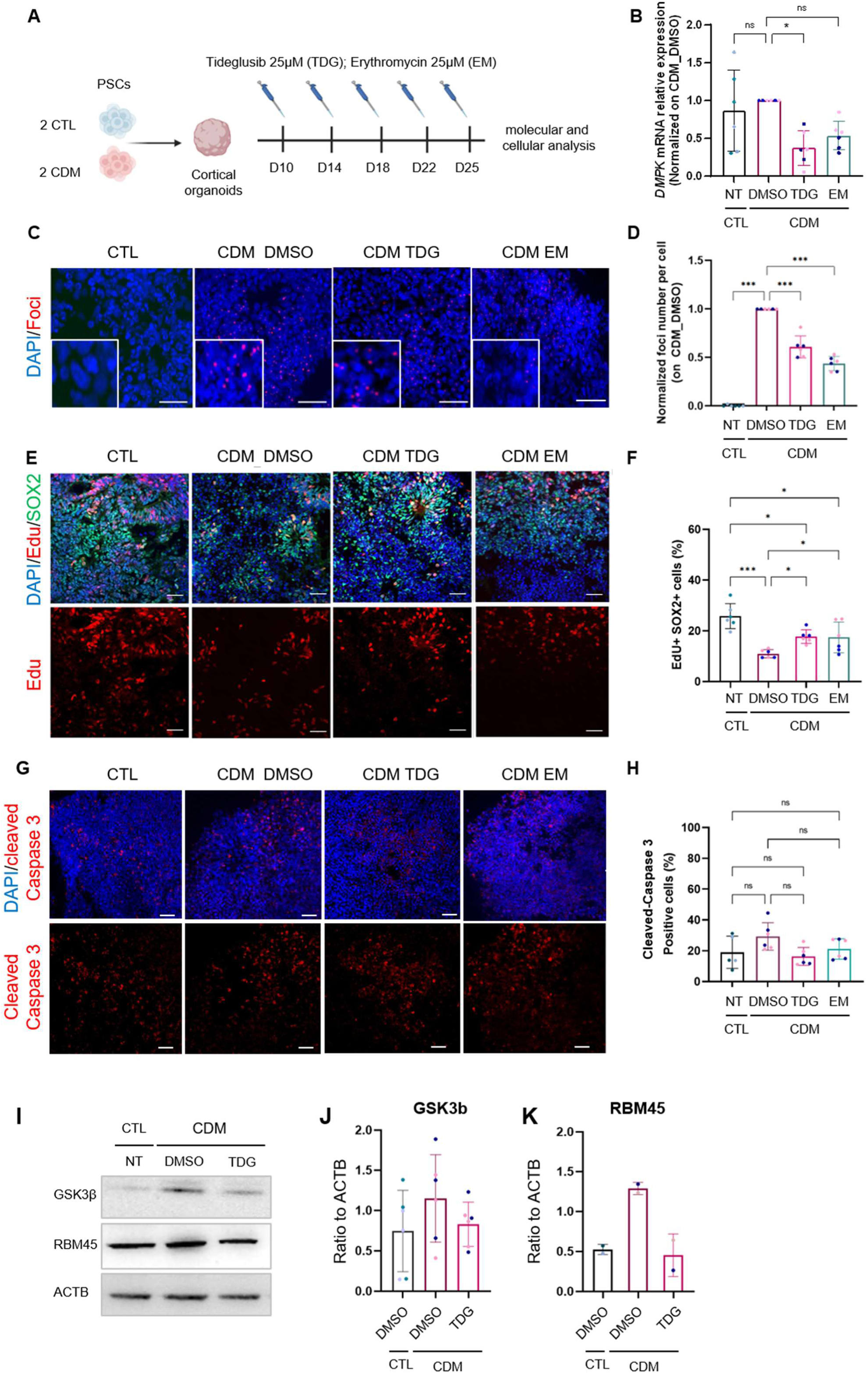
Small molecules treatment reduces CDM-associated molecular and cellular defects. **(A)** Scheme of the protocol used for organoids drug treatment. **(B)** RNA transcript levels of *DMPK* in 30-day-old organoids treated with or without tideglusib (TDG) and erythromycin (EM). **(C)** Representative images of RNA-FISH showing foci in 30-day-old CTL and CDM cortical organoids. **(D)** Quantification of DMPK foci number per cell by RNA FISH. (E-F) Representative images and quantification of EdU-labeled proliferating SOX2+ neural progenitor cells in control and CDM treated organoids. **(G-H)** Representative images and quantification of cleaved-caspase 3 staining in control, untreated and treated CDM organoids. **(I)** Western blot analysis showing protein level of GSK3β and RBM45 in control, untreated and treated CDM organoids. **(J-K)** Quantification of GSK3β and RBM45 levels as ratios to ACTB levels. Scale bars 50 µM. Colors indicate distinct cell lines within each genotype, and individual dots represent the mean value of three organoids from one independent differentiation experiment. Data are presented as mean ± SD. Statistical analysis was performed using one-way ANOVA followed by Dunnett’s multiple comparisons test. Statistical significance is indicated as follows: *p < 0.05; ***p < 0.001.

To confirm target engagement, GSK3β levels were measured after Tideglusib treatment. Elevated GSK3β protein in CDM organoids was normalized by treatment (**Figure 5I and J**). RBM45, an RNA-binding protein previously reported to be upregulated in DM1 and regulated by GSK3 activity, was also elevated in CDM organoids and restored to control levels following Tideglusib exposure (**Figure 5I and K**). These results demonstrate that pharmacological intervention in CDM cortical organoids reduces toxic RNA burden and partially rescues both molecular and cellular defects. The findings support the utility of human cortical organoids as a platform to evaluate therapeutic strategies targeting CNS manifestations of DM1.

## DISCUSSION

Congenital myotonic dystrophy type 1 (CDM) represents the most severe form of DM1 and is associated with profound CNS involvement. Despite increasing recognition of neurodevelopmental abnormalities in CDM patients, therapeutic strategies have predominantly focused on skeletal muscle pathology, leaving brain manifestations largely unexplored. Using human cortical organoids, we demonstrate that toxic RNA foci and MBNL protein sequestration emerge early in SOX2⁺ neural progenitors, leading to impaired progenitor proliferation and subsequent neuronal-glial imbalance. These phenotypes are faithfully recapitulated in MBNL- depleted organoids, supporting a central role for MBNL loss of function in early human corticogenesis.

Human cortical organoids provide a physiologically relevant platform to model early stages of human neurogenesis and cortical organization. In cortical organoids derived from CDM PSCs, pathogenic intranuclear RNA foci were detected in both progenitors and postmitotic neurons, indicating that DM1 affects multiple stages of cortical development. At early stages, most neural progenitors displayed RNA foci together with marked proliferation defects in the absence of increased apoptosis, suggesting impaired progenitor fitness rather than overt cell death. These observations are consistent with previous studies on DM1 neural stem cells and myogenic precursors, where impaired proliferation has been associated with enhanced autophagy, premature senescence and altered cell cycle regulation (**Denis et al., 2013; Bargiela et al., 2019; Song et al., 2020; Conte et al., 2023; Beffy et al., 2010; Bigot et al., 2009**). A formal demonstration of background-matched causality would require derivation of CRISPR-corrected isogenic CDM lines. Although three control and CDM hiPSC lines were used in this study, we therefore cannot fully exclude that a portion of the magnitude of cross-genotype differences reflects donor-specific genetic background.

At later stages, CDM organoids exhibited a marked reduction in CTIP2⁺ deep-layer and SATB2⁺ superficial-layer neurons, together with an expansion of GFAP⁺ and NFIA⁺ glial populations, consistent with a disrupted neurogenic-to-gliogenic switch (**Morelli et al., 2022; De Serres-Bérard et al., 2025**). Similar alterations have been reported in other CDM brain organoid models. In CDM forebrain organoids, De Serres-Bérard et al. attributed the reduction in mature SATB2⁺ and NeuN⁺ neurons to an impaired neuronal migration of outer radial glia (oRG) whereas Morelli et al. reported hyperactivity of 60-day-old DM1 cortical neurons and showed that inhibition with NMDA receptor antagonists mitigates neuronal loss. Together, these findings suggest that neuronal depletion in CDM likely results from converging pathogenic mechanisms involving impaired progenitor dynamics, defective neuronal migration and altered neuronal network activity.

In parallel, we observed increased gliogenesis in CDM cortical organoids, in agreement with previous report (**Morelli et al., 2022**). Given the early proliferation defects affecting neural progenitors in CDM cortical organoids, these findings raise the possibility that DM1 interferes with the proper timing and execution of the neurogenic-to-gliogenic transition (**Miller et al., 2007**). Because neurons and astrocytes arise from a common neuroepithelial progenitor pool, the presence of RNA foci in these progenitors, combined with altered proliferative capacity, may bias lineage progression toward glial amplification. Consistent with this hypothesis, we detected an increased number of NFIA⁺ cells, key regulators of astroglial lineage specification, in 120-day-old CDM cortical organoids. Functional analysis further confirmed MBNL loss of function as a major driver of these developmental defects. Both *MBNL2* KO and *MBNL* triple-knockout organoids reproduced the progenitor proliferation defects and lineage abnormalities observed in CDM organoids, consistent with single-cell transcriptomics showing high MBNL expression in cycling radial glia (**De Serres-Berard, 2025**). Previous work in DM1 myoblasts and satellite cells further supports MBNL1/2’s role in controlling proliferation through autophagy and mTOR regulation (**Song, 2020**; **Bargiela, 2019**). These observations raise an important question regarding the causal relationship between early progenitor proliferation defects and subsequent impairments in neuronal and glial differentiation. Transient degradation of expanded CTG repeats during early stages of CDM organoid differentiation using Cas13- based approaches, as previously described by **Morelli et al., 2022**, may help determine whether correction of RNA toxicity is sufficient to prevent later neurodevelopmental abnormalities. Altogether, the pronounced vulnerability of neural progenitor populations together with the early onset of CNS manifestations in CDM strongly argues for therapeutic intervention at the earliest stages of development. Our findings further highlight the need for human neurodevelopmental models capable of evaluating not only molecular biomarkers of DM1 pathology, but also the impact of candidate therapies on corticogenesis and neural lineage progression. CDM cortical organoids also provide a powerful platform for the evaluation of CNS-targeted therapies. Tideglusib, a GSK3β inhibitor with promising clinical outcomes in congenital DM1 (**Horrigan,2020**), reduced RNA foci, rescued progenitor proliferation, and normalized GSK3β and RBM45 levels. Notably, tideglusib has also been reported to counteract proliferation defects in neural progenitors within organoids carrying mutations in SPG11 (**Pozner, 2018**) confirming its potential to rescue proliferation defects in human cortical models. However, given the central role of GSK3β pathway in neural development, further studies will be required to evaluate the long-term effects of prolonged tideglusib exposure on corticogenesis and neural maturation.

Our findings further reinforce MBNL sequestration as a central pathogenic mechanism in CDM and support therapeutic approaches aimed at releasing MBNL proteins from toxic RNA aggregates. In this context, Nakamori et al. showed that erythromycin treatment, in vitro, in a pre-clinical model and in DM1 patients, corrects multiple DM-associated defects by displacing MBNL proteins from RNA foci (**Nakamori, 2015; Jenquin, 2019; Nakamori, 2024**). Consistent with these observations, our results indicate that, without altering *DMPK* expression, erythromycin treatment reduces intranuclear aggregates in NSCs, leading to enhanced cell proliferation. Combining organoid miniaturization, high-throughput imaging, and multiplexed phenotypic analysis could accelerate drug discovery for DM1 (**Narazaki, 2025; Hergenreder, 2024; Negraes, 2021; Trujillo et al., 2019**).

In summary, CDM profoundly impacts neural development from the earliest stages, with MBNL sequestration as a key driver. Cortical organoids offer a unique, human-relevant model to dissect disease mechanisms and assess both efficacy and safety of novel CNS-targeted therapies, bridging preclinical studies and clinical translation for DM1.

## MATERIALS AND METHODS

### Generation of human cortical organoids

Cortical organoids were generated as previously described (**Paşca et al., 2015**) from two control and two CDM iPSC lines (**Table S1)** (**Tahraoui-Bories et al., 2023**) and one control and one CDM hES cell lines (**Yanovsky-Dagan et al., 2015**). Two additional iPSC lines with *MBNL2* or *MBNL1*/2/3 genes knock-out were derived from one of the two control iPSC lines (**Merien et al., 2021**). Briefly, iPSC and hES colonies were cultured and expanded with StemMACS^TM^ iPS Brew XF (Miltenyi Biotec®) medium for four days. Then, PSC lines were dissociated with Trypsin-EDTA 0,25% (Gibco^TM^). Suspended cells were subsequently transferred into a 96-well microplate (ThermoFisher®) in neural medium containing Neurobasal (Life Technologies), B- 27 supplement without vitamin A (Life technologies®), N2 supplement (Life Technologies®), DMEM-F12-glutamax (Life technologies®), 50mM β-mercapto-ethanol (Life Technologies®) and 10 000 U/mL penicillin and streptomycin (Gibco^TM^). For the first 24 h (day 0), the neural medium was supplemented with the ROCK inhibitor Y-27632 (EMD Chemicals). For neural induction, 200ng/ml Noggin (Peprotech®) and 10µM SB-431542 (R&D Systems) were added to the neural medium for the first six days. On day 7 of differentiation, the floating spheroids were embedded into Matrigel (Corning®) drops and transferred into ultra-low attachment 6- well plates. For the following eighteen days, the medium was supplemented with 20 ng/ml FGF2 (Miltenyi Biotec®) and 20 ng/ml EGF (R&D Systems®). To promote differentiation of the neural progenitors into neurons, FGF2 and EGF were replaced with 10 ng/ml BDNF (Peprotech®) and 20 ng/ml NT3 (Peprotech®) starting at day 25, while from day 43 onwards only neural medium without growth factors was used for medium changes every two days. Organoids cultures were kept in suspension under rotation (95 rpm).

### Cortical organoid treatment

Human cortical organoids were treated with 25µM tideglusib (Sigma Aldrich®) and 25µM erythromycin (MedChem Express®) starting from day 10 of differentiation until day 25. Compounds were freshly diluted in the neural medium supplemented with 20 ng/ml FGF2 (Miltenyi Biotec®) and 20 ng/ml EGF (R&D Systems®). CDM organoids received an equivalent concentration of the vehicle (0.1% DMSO). The culture medium containing the respective compounds or vehicle was replaced every four days to ensure continuous exposure. At day 25, organoids were collected for downstream analysis.

### RT-PCR and Agilent DNA chips analysis for Alternative Splicing

Total RNAs were isolated using the RNAeasy Mini extraction kit (Qiagen®) from human cortical organoids according to the manufacturer’s protocol. RNA levels and quality were checked using NanoDrop® technology. RNA was reverse transcribed using random hexamers and the Superscript III Reverse Transcriptase kit (Invitrogen®) according to the manufacturer’s protocol. For splicing analysis, PCR amplification was carried out with recombinant Herculase II Fusion DNA polymerase (Agilent®) and primers listed in **Table S1**. The amplification was performed using a first step at 94°C for 3 minutes followed by 35 cycles of 30 sec at 94°C, 30 sec at 60°C, 1 min at 72°C, and a final 10 min extension at 72°C. The PCR products were analyzed using Agilent® DNA chips and quantified with the BioAnalyzer 2100. The percent spliced-in (PSI) value corresponds to the fraction of mRNAs that contain an exon and was calculated as the ratio of the density of the exon inclusion band to the sum of the densities of inclusion and exclusion bands, expressed as a percentage.

### Gene expression analysis by quantitative PCR

Total RNA was extracted using the RNeasy Mini kit (Qiagen®) and was reverse transcribed using random hexamers and the Superscript III Reverse Transcriptase kit (Invitrogen®). Quantitative PCR reactions were carried out in 384-well plates using a QuantStudio 12K Flex Real-Time PCR System (Applied Biosystems®) with Power SYBR Green 2× Master Mix (Life Technologies®), 2.5 μl of cDNA and primers (**Table S1**) in a final volume of 10 μL. All analyses were performed with at least three technical replicates per plate. The 2^-ΔΔ^ Ct method was used to determine the relative expression level of each gene. Expression data were normalized to *18S*.

### Protein extraction and western blot analysis

Western blot analyses were performed as previously described (**Denis et al., 2013**). Briefly, human cortical organoids were lyzed in RIPA 1X buffer (Sigma-Aldrich ®) containing protease inhibitors (Sigma-Aldrich ®) and phosphatase inhibitors (Roche®). Proteins were quantified by Pierce BCA Protein Assay kit (Pierce®) using a multiplate colorimetric reader, CLARIOstar (BMG Labtech®). Protein extracts (10-20 μg) were loaded on a 4%–12% SDS-PAGE gradient gel (NuPage Bis-Tris gels, Invitrogen®) and transferred onto Gel Transfer Stacks Nitrocellulose membranes (Invitrogen®) using the iBlot2 Dry Blotting System (Invitrogen®). After blocking with 5% non-fat milk for 1 hour at room temperature, membranes were incubated overnight at 4°C with the following primary antibodies: mouse anti-MBNL2 (Santa Cruz®, sc-136167 3B4, 1:1000), rabbit anti-GSK3β (Cell Signaling Technology ®, 1 :1000), rabbit anti-RMB45 (Abcam ®, 1 :1000), rabbit anti-Histone H3 (Cell Signaling Technology ®, 1 :1000), rabbit anti-Phospho-Histone H3 (Cell Signaling Technology®, 1 :1000) and rabbit or mouse anti ACTB (LICORbio®, 1:1000). Membranes were then incubated with HRP-conjugated secondary antibody (1:10000), and immunoreactive bands were revealed using Amersham ECL Prime Western Blotting Detection Reagents (Cytiva Life Science®). Signals were visualized using ImageQuant software and quantified using ImageJ.

### Cryopreservation of cortical organoids

Cortical organoids were fixed with 4% paraformaldehyde for 30 min at room temperature, transferred to 30% sucrose for 24-48 hours at 4°C. Organoids were transferred into embedding reagent O.C.T (Sakura®) snap-frozen on dry ice and stored at −80 °C. For immunofluorescence, 15 µm thick sections were performed using cryostat and transferred onto Superfrost^TM^ slides (Epredia^TM^).

### Immunostaining

Cryosections were incubated in 1% BSA, 0.1% Triton in PBS for 1 hour at room temperature and then with primary antibodies overnight in 4°C. Primary antibodies used in this study were: rat anti-CTIP2 (Abcam®, 1:250), mouse anti-TUJ1 (Biolegend®, 1:500), rabbit anti-SOX2, (EMD Mllipore®, 1:500), chicken anti-GFAP (Abcam®, 1:500), mouse anti-S100β (Sigma Aldrich ®, 1:500), mouse anti-HuC/D neuronal protein (Invitrogen®, 1:250), rabbit anti-cleaved caspase 3 (Cell Signaling Technology®, 1:600) and rabbit anti-TBR1 (Abcam®, 1:250) mouse anti-SATB2 (Abcam, 1:250), mouse anti-MBNL1 (Abcam®, 1:100), mouse anti- MBNL2 (SantaCruz®, 1:100), rabbit anti-PAX6 (Biolegend®, 1:500), rabbit anti-NFIA (Novus®, 1:200). The sections were incubated with secondary antibodies (Alexa Fluor 488-, 555- and 647-conjugated antibodies, Life Technologies®, 1:1000) for 2 hours at room temperature. Nuclei were stained using DAPI solution (1 mg/mL). The slides were mounted using Fluoromont (ThermoFisher®) reagent. Images were acquired using Zeiss Spinning Disk microscope with the 20X objective. All cells expressing a specific marker were counted on at least 3 sections per organoid and normalized to the total number of cells using ImageJ software.

### Fluorescent In Situ Hybridization

Cryosections were incubated 30 min in 70% Ethanol (VWR®) at room temperature. After three washes in PBS, sections were rehydrated with a solution of 5mM of MgCl2 (Sigma-Aldrich®) for 10 min at room temperature and then sequentially incubated with prehybridization buffer (50mM Phosphate buffer, 40% formamide, 2X SSC) for 10 min and hybridization buffer (Prehybrydization buffer with 0,2% BSA and 7% Dextran, 50µl/well) containing 300ng/mL of the (CAG)10-Cy5 probe (5’Alexa 647-TTCTTATTCTTCAgCAgCAgCAgCAgCAgCAgCAgCAgCAg3’) (Operon) overnight at 37°C. Washing steps consisted in pre-warmed washing buffer (Prehybrydization buffer + 0.2% BSA) for 1 hour at 37°C. Finally, cryosections were washed twice in PBS buffer and incubated 10 min in a Hoechst 33528 solution (Sigma-Aldrich®). Images were acquired using Zeiss Spinning Disk microscope with the 20X objective. Nuclei and nuclear foci were detected using QuPath, and quantitative analysis was subsequently performed with ImageJ software.

### Proliferation analysis

Proliferation was assessed with EdU staining conducted using Click-iT™ EdU imaging kit (Invitrogen®, Carlsbad, CA) according to the manufacturer’s protocol. Briefly, cortical organoids were incubated with 20 µM EdU in culture medium. After 4 hours of incubation, organoids were immediately fixed in 4% paraformaldehyde for 30 min and then permeabilized with 1% BSA, 0.1% Triton in PBS at room temperature. Organoids were then incubated in 30% sucrose for 24-48 hours. Organoids were transferred into embedding reagent O.C.T and cryosectioned using cryostat. Slides were incubated with a Click-iT™ reaction cocktail containing Click-iT™ reaction buffer, CuSO_4_, Alexa Fluor® 555 Azide, and reaction buffer additive for 45 min while protected from light. The sections were washed once in PBS. Additional label was performed with primary antibody rabbit anti-SOX2, (EMD Mllipore®, 1:500). The sections were washed with PBS and incubated with secondary antibody Alexa Fluor 488 and DAPI solution for 2 hours. Slides were washed with PBS and mounted using Fluoromont (Thermo Fisher®) reagent.

### Statistical analysis

All data were processed using GraphPad Prism 8®. Values are presented as mean ± SD. For comparisons involving two independent groups, statistical analyses were performed using an unpaired two-tailed Student’s t-test. When more than two independent groups were compared, statistical analyses were performed using ordinary one-way analysis ANOVA followed by Dunnett’s multiple comparisons test, with CTL as the control group. Non-parametric analysis were performed using a Kruskal–Wallis test followed by Dunn’s multiple comparisons test, comparing each group with CTL as the control group. For analysis involving two factors, genotype and time, statistical analyses were performed using ordinary two-way ANOVA followed by Dunnett’s multiple comparisons test, with CTL as the control group at each time point. Statistical significance was set at p < 0.05. Values were considered significant as follows: *p < 0.05; **p < 0.01; ***p < 0.001; ****p < 0.0001.

## LIST OF SUPPLEMENTARY MATERIALS

Supplementary Figures 1 to 3 and Supplementary Table1.

## Acknowledgments

We thank Noémie Beranger-Currias and Sandra Pourtoy for their technical help. I-Stem is part of the Biotherapies Institute for Rare Diseases (BIRD) and is financially supported by the Association Française contre les Myopathies (AFM-Téléthon). This work was in part conducted at the stemCARE platform of I-Stem supported by the French bioclusters Genopole and Genother (Evry, France) and GIS IBISA. This work benefited from a government grant managed by the Agence Nationale de la Recherche under the France 2030 program 23- BIOC-0003-04.

## Author contributions

**Conceptualization**: A.A., M.G.P., S.B. and C.M.; **Methodology**: A.A., S.B. and C.M.; **Software**: J.P.; **Validation**: A.A., S.B. and C.M.; **Formal analysis**: A.A.; **Investigation**: A.A., M.B., A.B., H.M., L.E.K., S.T. and S.B.; **Resources**: J.P., M.B., A.B., O.C., L.C., A.B.; **Data curation**: A.A.; **Writing – original draft**: A.A., S.B. and C.M.; **Writing – review & editing**: M.G.P, G.G., S.B. and C.M.; **Visualization**: A.A. and S.B.; **Supervision**: C.M.; **Project administration**: C.M.; **Funding acquisition**: M.G.P. and C.M.

## Data and materials availability

Materials can be asked for availability to Cécile Martinat

