## Supplemental figure 1 to 3 and Supplementary Table 1 for "Cortical organoids from congenital DM1 PSCs reveal MBNL-dependent corticogenesis defects and enable preclinical testing of therapeutic compounds"

### Supplementary materials

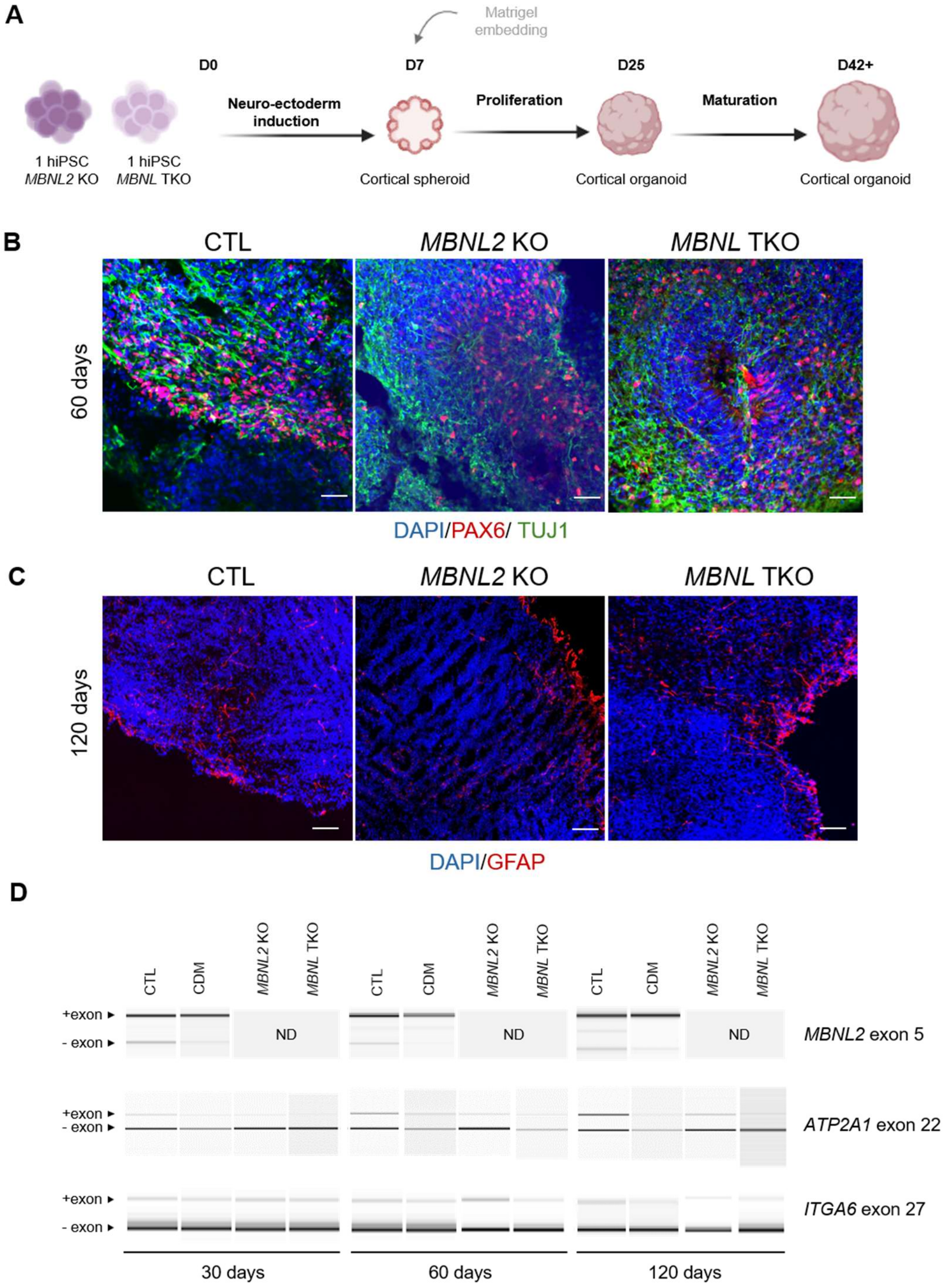


**Supplementary Figure 1.** Generation of cortical organoids derived from *MBNL2*, *MBNL1/2/3* KO and control pluripotent stem cells. **(A)** Schematic representation of the protocol used to generate control and MBNL-depleted cortical organoids. **(B-C)** Immunostaining of PAX6, TUJ1 and GFAP markers in control and MBNL-depleted cortical organoids at days 60 and 120 of differentiation. Scale bars, 50 µm. **(D)** Representative images of *MBNL2*, *ATP2A1* and *ITGA6* alternative splicing analyzed by RT-PCR in 30-, 60- and 120-day-old cortical organoids. ND: not detected.

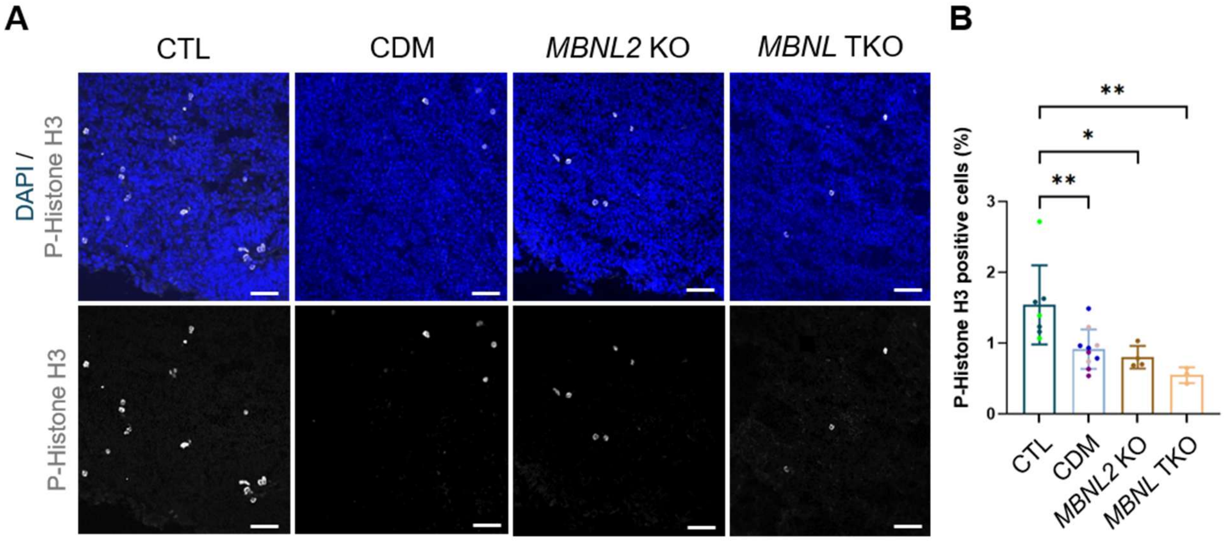


**Supplementary Figure 2. Impaired proliferation in CDM and *MBNLs* KO cortical organoid. (A)** Representative images of phospho-histone H3 cells in 30-day-old cortical organoids. **(B)** Quantifications of phospho-histone H3 cells. Colors indicate distinct cell lines within each genotype, and individual dots represent the mean value of three organoids from one independent differentiation experiment. Data are presented as mean ± SD. Statistical analysis was performed using two-way ANOVA followed by Dunnett's multiple comparisons test. Statistical significance is indicated as follows: *$p < 0.05$; **$p < 0.01$.

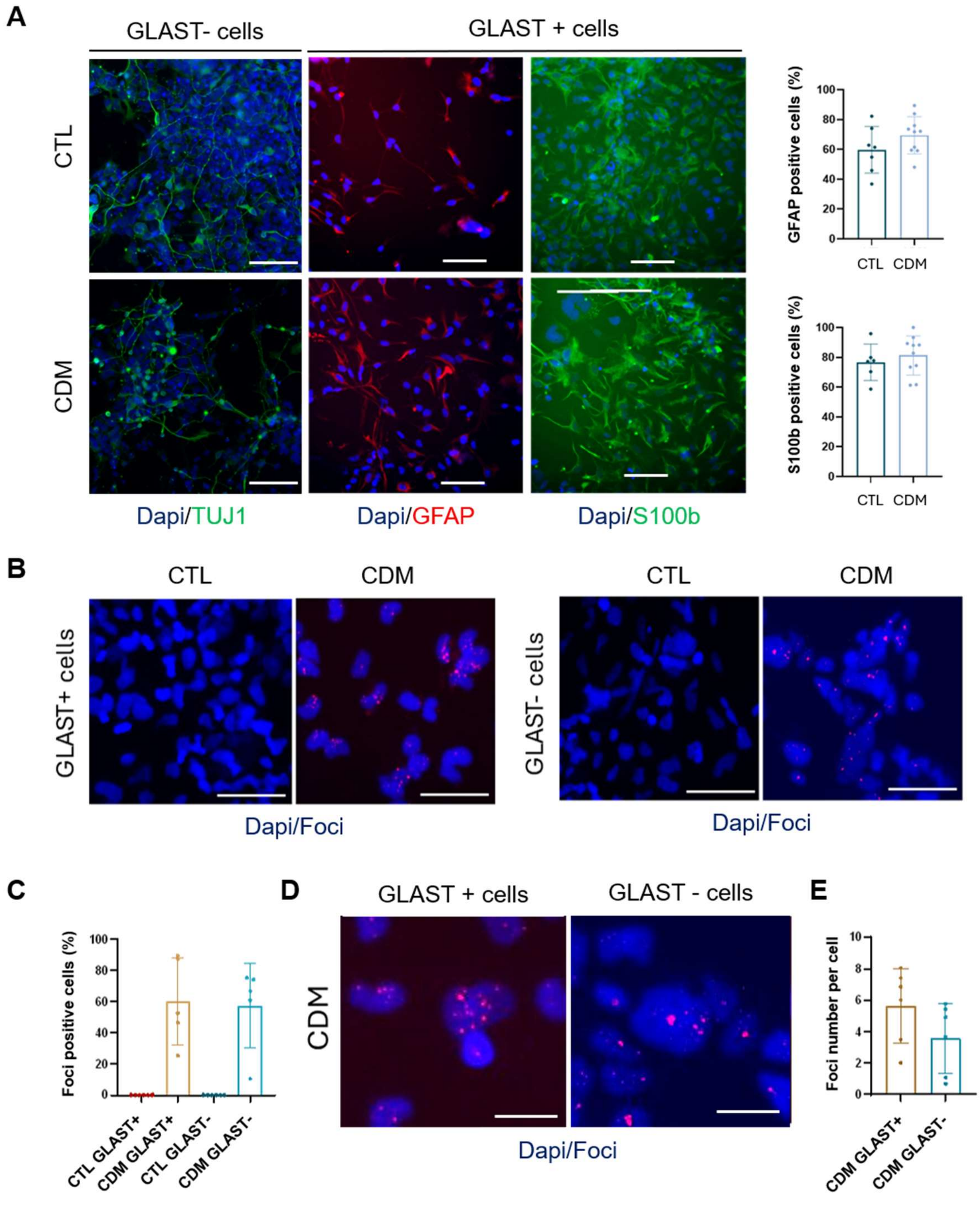


**Supplementary Figure 3. Characterization of neural and glial cells derived from dissociated CTL and CDM cortical organoids.** (**A**) Representative images and quantification of TUJ1, GFAP and S100β positive cells detected by immunostaining in GLAST$^+$ and GLAST$^-$ cellular populations purified from 120-day-old CTL and CDM cortical organoids. Scale bars, 50 μm. (**B-E**) Representative images and quantification of RNA FISH showing DMPK foci in GLAST$^+$ and GLAST$^-$ cells purified from 120-day-old CTL and CDM cortical organoids. Scale bars, 10 μm. Data are presented as mean ± SD from two independent experiments performed using one CTL hiPSC line and two DM1 hiPSC lines. For each independent experiment and each cell line, 3–4 technical replicate wells were analyzed.

**Supplementary Table 1**

**Characterization of the CTG repeat length in the three DM1-mutated pluripotent stem cell lines used in the study using PacBio long-read sequencing**

| Cell line | Genotype | Median size of CTG repeats |
|---|---|---|
| **iPSC 61c7** | DM1 | 5 and 3358 |
| **iPSC 62c12** | DM1 | 12 and 2508 |
| **hES SZ-DM6** | DM1 | 11 and 1046 |

**Primers used for alternative splicing analysis**

| | | |
|---|---|---|
| ATP2A1 exon 22 | Forward | AGTTCGTTGCTCGGAACTACC |
| | Reverse | GCCTGAAGATGTGTCACTATCG |
| ITGA6 exon 27 | Forward | GAGTGACTGTGTTTCCCTCAAAGAC |
| | Reverse | CAGCCACGCCAAAAATAAAGG |
| MBNL2 exon 5 | Forward | AGGCCAAAATCAAAGCTGCG |
| | Reverse | GTGAGAGCCTGCTGGTAGTG |

**Primers used for qPCR analysis**

| | | |
|---|---|---|
| DMPK | Forward | CGGGCCGTCCGTGTT |
| | Reverse | CCAGAGCAGGGCGTCATG |
| MBNL1 | Forward | CTCAGTCGGCTGTCAAATCA |
| | Reverse | ACTGGTGGGAGAAATGCTG |
| MBNL2 | Forward | CTTCATCCAGTGCCCACTTT |
| | Reverse | TGGAACTCCCTGCATAACTC |
| 18S | Forward | GAGGATGAGGTGGAACGTGT |
| | Reverse | TCTTCAGTCGCTCCAGGTCT |
